# RNA sensing via LGP2 is essential for the induction of a type I IFN response in ADAR1 deficiency

**DOI:** 10.1101/2021.10.27.465188

**Authors:** Jorn E. Stok, Timo Oosenbrug, Laurens R. ter Haar, Dennis Gravekamp, Christian P. Bromley, Santiago Zelenay, Caetano Reis e Sousa, Annemarthe G. van der Veen

**Author notes:** These authors contributed equally to this work (Jorn E. Stok, Timo Oosenbrug).

## Abstract

RNA editing by the enzyme Adenosine Deaminase Acting on RNA 1 (ADAR1) is an important mechanism by which cells avoid innate immune responses to some endogenous RNAs. In ADAR1-deficient cells, unedited self RNAs can form base-paired structures that resemble viral RNAs and inadvertently activate antiviral innate immune pathways that lead to the induction of type I interferon (IFN). Rare mutations in ADAR1 cause Aicardi-Goutières Syndrome (AGS), a severe childhood autoinflammatory syndrome that is characterized by chronic and excessive type I IFN production and developmental delay. Conversely, ADAR1 dysfunction and consequent type I IFN production helps restrict tumor growth and potentiates the activity of some chemotherapy drugs. Induction of type I IFN in ADAR1-deficient cells is thought to be due to triggering of the cytosolic RIG-I-like receptor (RLR), MDA5, by unedited self RNAs. Here, we show that another RLR, LGP2, also has an essential role. We demonstrate that ADAR1-deficient human cells fail to mount a type I IFN response in the absence of LGP2 and this involves the canonical function of LGP2 as an RNA sensor and facilitator of MDA5-dependent signaling. Further, we show that the sensitivity of tumor cells to ADAR1 loss requires the presence of LGP2. Finally, we find that type I IFN induction in tumor cells depleted of ADAR1 and treated with some chemotherapeutics is fully dependent on the expression of LGP2. These findings highlight a central role for LGP2 in self RNA sensing with important clinical implications for the treatment of AGS as well as for the potential application of ADAR1-directed anti-tumor therapy.

## Introduction

Receptors of the innate immune system continuously sample the intra- and extracellular environment for signs of an ongoing infection. Viral infections can be detected through the presence of viral nucleic acids in the cytosol of infected cells (Rehwinkel & Gack, 2020; Goubau *et al*, 2013). Upon encountering viral DNA or RNA, cytosolic nucleic acid sensors, most notably cGAS or RIG-I-like receptors (RLRs), respectively, initiate an antiviral type I interferon (IFN) response (Ablasser & Hur, 2020; Rehwinkel & Gack, 2020). While a type I IFN response is important for defense against viral infections, its inadvertent activation by self-derived nucleic acids induces a sterile inflammatory response that causes immunopathology (Schlee & Hartmann, 2016). Cellular mechanisms that ensure the discrimination between foreign and endogenous nucleic acids are therefore critical to avoid autoinflammation (Schlee & Hartmann, 2016).

RNA modification by the enzyme Adenosine Deaminase Acting on RNA (ADAR1) constitutes an important mechanism by which cells ensure self/non-self RNA discrimination (Heraud-Farlow & Walkley, 2016; Quin *et al*, 2021; Uggenti *et al*, 2019). Through modification of endogenous RNA, ADAR1 prevents the activation of cytosolic RNA sensors, including RLRs, by cellular RNA molecules and the unwanted induction of an antiviral type I IFN response (Heraud-Farlow & Walkley, 2016; Uggenti *et al*, 2019; Quin *et al*, 2021). The importance of ADAR1 is highlighted by the severe consequences of *ADAR1* mutations in patients with Aicardi-Goutières Syndrome (AGS) (Rice *et al*, 2012; Rodero & Crow, 2016). This rare genetic disorder belongs to the spectrum of type I interferonopathies, which are characterized by the constitutive induction of an antiviral type I IFN response in the absence of an infection (Rodero & Crow, 2016). The autoinflammatory condition that arises from inherited *ADAR1* mutations leads to severe (neuro)pathological features (Rice *et al*, 2017; Livingston & Crow, 2016). Notably, ADAR1 has also emerged as an attractive target for novel immunotherapeutic approaches in cancer (Bhate *et al*, 2019). A subset of tumor cells is sensitive to growth arrest upon knockdown or knockout of ADAR1, both *in vivo* and *in vitro* (Ishizuka *et al*, 2019; Gannon *et al*, 2018; Liu *et al*, 2019). In addition, intra-tumoral loss of ADAR1 increases sensitivity to treatment with immune checkpoint inhibitors and overcomes resistance to such inhibitors *in vivo* (Ishizuka *et al*, 2019). Finally, depletion of ADAR1 in cancer cells potentiates the efficacy of epigenetic therapy and increases type I IFN induction (Mehdipour *et al*, 2020). Understanding the precise mechanism by which ADAR1 (dys)function impacts on innate immunity is therefore essential to better understand its disease-causing role in interferonopathies as well as its therapeutic potential in cancer.

ADAR1 exists as two isoforms. The nuclear p110 isoform is constitutively expressed, while the p150 isoform is induced by type I IFN receptor signaling and resides primarily in the cytoplasm (Heraud-Farlow & Walkley, 2016; Quin *et al*, 2021). Both isoforms act on base-paired RNA to deaminate adenosines and convert them to inosines. A-to-I editing is amongst the most widespread base modifications in mammals. Besides site-specific A-to-I editing, which can alter open reading frames, miRNA seed sequences or RNA splice sites, there is also highly promiscuous and abundant editing of base-paired RNAs with long regions of high complementarity such as transcripts spanning inverted repeat Alu (IR-Alu) elements (Eisenberg & Levanon, 2018). Without editing, such base-paired structures would resemble double-stranded RNAs (dsRNAs) that are abundantly found in cells infected with some viruses. Unedited self RNA molecules are therefore prone to activate antiviral innate immune mechanisms, such as protein kinase R (PKR) (Chung *et al*, 2018), OAS1/RNase L (Li *et al*, 2017) and the RLR pathway (Liddicoat *et al*, 2015; Pestal *et al*, 2015; Mannion *et al*, 2014). While activation of PKR and OAS/RNase L causes translational shutdown and cell death, RLR engagement initiates the type I IFN response.

The link between ADAR1 editing and RLR activation was first demonstrated in a series of mouse studies. In mice, genetic loss of ADAR1 p110 and p150, p150 alone, or knock-in of an editing-deficient ADAR1 mutant (*Adar ^E861A/E861A^*) results in embryonic lethality, fetal liver disintegration, hematopoiesis defects, and an elevated type I IFN signature (Wang *et al*, 2004; Hartner *et al*, 2004; Ward *et al*, 2011; Liddicoat *et al*, 2015). The embryonic lethality of ADAR1 null or editing-deficient mice can be rescued by the concurrent deletion of the RLR family member MDA5 (melanoma differentiation-associated gene 5) or the downstream signaling hub MAVS (mitochondrial antiviral signaling, also known as VISA, Cardif, IPS-1), but not another RLR, RIG-I (retinoic-acid-inducible gene I) (Liddicoat *et al*, 2015; Mannion *et al*, 2014; Pestal *et al*, 2015; Heraud-Farlow & Walkley, 2016). In addition, loss of MDA5 or MAVS also eliminates the type I IFN signature in these mice. These observations indicate that unedited RNA mediates its immunostimulatory effects via MDA5 and MAVS and that the type I IFN response plays an important role in the immunopathology caused by loss of ADAR1.

MDA5 normally detects RNA from certain viral species, such as *Picornaviridae* (Dias Junior *et al*, 2019). It senses long stretches of dsRNA or base-paired single stranded RNA, on which it oligomerizes to form filamentous structures (Dias Junior *et al*, 2019; Rehwinkel & Gack, 2020). In contrast, RIG-I is activated by 5’ di- or triphosphate moieties at the base-paired extremities of certain viral RNA species (Goubau *et al*, 2013; Rehwinkel & Gack, 2020). Activation of RIG-I or MDA5 by their respective RNA substrates leads to conformational changes that allow their N-terminal CARD domains to interact with the CARD domains of the adaptor MAVS (Sohn & Hur, 2016). This, in turn, leads to MAVS activation and subsequent phosphorylation and activation of the transcription factors IRF3 and NF-*κ*B, which mediate the transcription of type I IFNs (most notably IFN-α subtypes and IFN-β), type III IFNs and other pro-inflammatory cytokines (Goubau *et al*, 2013; Rehwinkel & Gack, 2020). Upon secretion, type I IFNs activate the IFN receptor (IFNAR) and induce JAK-STAT signaling, which results in the transcriptional upregulation of hundreds of IFN-stimulated genes (ISGs), which establish an antiviral state (Schoggins *et al*, 2011; Schneider *et al*, 2014). LGP2 (laboratory of genetics and physiology 2) is the third and least well-understood member of the RLR family. LGP2 lacks the N-terminal CARD domains and is therefore not able to signal via MAVS (Rodriguez *et al*, 2014; Rehwinkel & Gack, 2020). Instead, LGP2 modulates the function of RIG-I and MDA5 during viral infection. While LGP2 suppresses RIG-I signaling, it synergizes with MDA5 to potentiate the sensing of certain RNA viruses (Rodriguez *et al*, 2014; Rehwinkel & Gack, 2020). Akin to MDA5-deficient mice, LGP2 knockout mice display increased sensitivity to infection with encephalomyocarditis virus (EMCV), a member of the *Picornaviridae* family (Satoh *et al*, 2010; Venkataraman *et al*, 2007). Mechanistically, LGP2 is incorporated into MDA5 filaments and enhances the interaction between MDA5 and RNA, thereby increasing the rate of MDA5 filament formation (Duic *et al*, 2020; Bruns *et al*, 2014). Simultaneously, LGP2 enhances the dissociation of MDA5 filaments in an ATP-dependent manner and generates shorter filaments that have greater agonistic activity than longer filaments (Duic *et al*, 2020; Bruns *et al*, 2014). Structural studies demonstrated that LGP2 primarily binds the ends of dsRNA, although it can also coat dsRNA in a similar fashion as MDA5 (Uchikawa *et al*, 2016). Thus, LGP2 promotes rapid MDA5-dsRNA filament formation yet yields shorter filaments, ultimately leading to enhanced downstream signaling and an increased type I IFN response.

LGP2 also impacts on type I IFN responses through alternative routes that are independent from its role as typical RNA sensor. Wild-type LGP2, as well as mutants that fail to hydrolyze ATP or bind RNA, interact with MAVS at steady state and block the interaction between RIG-I and MAVS, thereby limiting RIG-I-mediated MAVS activation (Esser-Nobis *et al*, 2020). Upon stimulation with the dsRNA mimic poly(I:C), LGP2 releases MAVS for interaction with RIG-I (Esser-Nobis *et al*, 2020). LGP2 additionally limits RIG-I signaling and potentiates MDA5 signaling by a direct protein-protein interaction with the dsRNA-binding protein PACT (Sanchez David *et al*, 2019). Furthermore, LGP2 inhibits Dicer-mediated processing of dsRNA (Van der Veen *et al*, 2018), perhaps to preserve dsRNA substrates for the full-blown activation of the type I IFN response. Conversely, LGP2 may negatively regulate the antiviral type I IFN response by associating and interfering with the function of TRAF ubiquitin ligases, in a manner that is independent of ATP hydrolysis or RNA-binding (Parisien *et al*, 2018). Finally, LGP2 controls CD8^+^ T cell survival and fitness during West Nile virus and lymphocytic choriomeningitis virus infection in mice, pointing to cell-type specific functions (Suthar *et al*, 2012).

Both MDA5 and RIG-I can bind and be activated by endogenous RNA in various contexts (Stok *et al*, 2020; Dias Junior *et al*, 2019; Streicher & Jouvenet, 2019). LGP2 has predominantly been studied upon viral infection or mimics thereof. A recent study demonstrated that mice bearing a mutation in ADAR1 that abolishes binding to dsRNA in its unusual Z-conformation (Z-RNA) suffer from postnatal growth retardation and mortality and have a mild type I IFN signature, which can be reverted by crossing these mice with MDA5, MAVS, PKR, as well as LGP2 knockout mice (Maurano *et al*, 2021). The extent to which LGP2 is required for type I IFN induction in response to unedited RNA species more broadly (aside from Z-RNA), the molecular mechanism that is involved, and whether it is required in humans is unclear.

Here, we investigated the role of human LGP2 in induction of type I IFNs caused by ADAR1 deficiency. Using various genetic approaches and model systems, we demonstrate that LGP2 is essential for this induction, in a manner that involves its classical function as RNA sensor. Importantly, we further demonstrate that LGP2 is required both for sensing of unedited RNA as well as for reduced cell growth upon loss of ADAR1 in tumor cells. Finally, treatment of ADAR1-depleted tumor cells with epigenetic repressors, a promising strategy for cancer therapy, potentiates the type I IFN response in an LGP2-dependent manner. Our findings provide molecular insight into the effector mechanisms that are engaged upon dysregulation of ADAR1, with important clinical implications for the field of interferonopathies as well as cancer.

## Results

### Human LGP2 is required for the induction of a type I IFN response upon depletion of ADAR1

To investigate the role of human RLRs in the induction of type I IFN caused by absence of ADAR1, we first knocked out RIG-I, MDA5, or LGP2 in the human monocytic leukemia cell line THP-1 using CRISPR/Cas9-mediated genome engineering. Correct gene ablation was confirmed by immunoblotting cells treated with recombinant type I IFN to upregulate the expression of RIG-I, MDA5, and LGP2, which are encoded by ISGs themselves (Fig. 1A). Intact type I IFN receptor signaling was verified by monitoring ISG60 upregulation (Fig. 1A). For each RLR, two knockout clones were differentiated into macrophages and transfected with siRNAs targeting both isoforms of ADAR1. Despite the modest efficiency of the knockdown, we observed a clear upregulation of transcripts encoding IFN-β and the ISG IFIT1 in wild-type cells, indicative of type I IFN induction (Fig. 1B). Notably, loss of LGP2 completely abrogated type I IFN induction and signaling upon ADAR1 depletion (Fig. 1B). Loss of MDA5, but not RIG-I, also interfered with the type I IFN response, consistent with published literature (Heraud-Farlow & Walkley, 2016). Note that throughout the manuscript ADAR1 knockdown efficiency is monitored through measurement of p110 expression levels, as analysis of the p150 isoform leads to distorted results due to its IFN-inducible nature. Together these data indicate that, besides MDA5, expression of LGP2 is crucial for the induction of a type I IFN response in ADAR1 deficiency.

**Figure 1:**
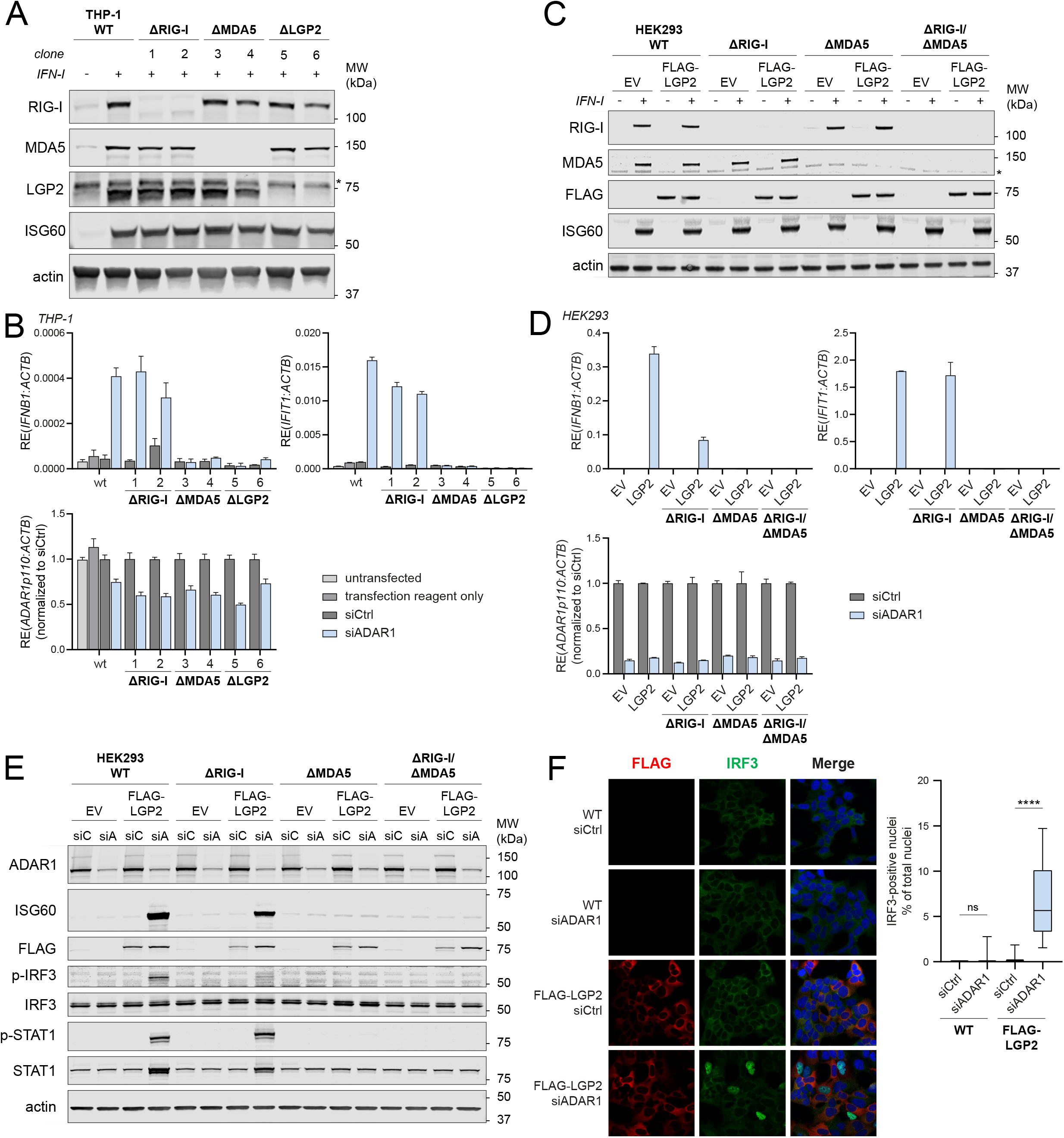
Human LGP2 is essential for the induction of a type I IFN response upon depletion of ADAR1. **A)** THP-1 monocytes were genetically engineered to knockout RIG-I, MDA5, or LGP2 using CRISPR/Cas9. Cells were differentiated towards macrophage-like cells using PMA and treated with recombinant type I IFN to upregulate RLR expression. Correct gene editing and intact type I IFN responsiveness was validated by SDS-PAGE and immunoblotting using the indicated antibodies (n=3). *, non-specific band. **B)** Cells generated in (A) were differentiated using PMA and transfected with a control siRNA (siCtrl) or an ADAR1-targeting siRNA (siADAR1). The type I IFN response was monitored 56h post-transfection by RT-qPCR analysis to determine IFN-β and IFIT1 transcript expression, normalized to a housekeeping gene (ACTB). ADAR1 knockdown efficiency was monitored by ADAR1 p110 expression, normalized to ACTB and displayed relative to siCtrl. Data from a representative experiment are shown with mean ± s.d. (n=3). **C)** HEK293 cells were genetically engineered to knockout RIG-I, MDA5, or both, and subsequently subjected to retroviral transduction to stably express FLAG-LGP2 or an empty vector (EV). Correct gene editing and intact type I IFN responsiveness was validated by SDS-PAGE and immunoblotting using the indicated antibodies (n=2). *, non-specific band. **D)** Cells generated in (C) were transfected with siCtrl or siADAR1. The type I IFN response and ADAR1 knockdown efficiency was monitored 78h post-transfection as in (B). Data from a representative experiment are shown with mean ± s.d. (n=4). **E)** Cells generated in (C) were transfected with siCtrl (siC) or siADAR1 (siA). Protein lysates were prepared 78h post-transfection, followed by SDS-PAGE and immunoblotting using the indicated antibodies (n=2). **F)** HEK293 cells and FLAG-LGP2-expressing HEK293 cells were transfected with siCtrl or siADAR1 and subsequently plated on coverslips for immunofluorescence. Cells were fixed, permeabilized and stained 72h post-transfection with anti-FLAG and anti-IRF3 antibodies. Total nuclei (>450 nuclei per experimental condition) and IRF3-positive nuclei were counted using semi-automated software analysis and plotted as percentage IRF3-positive nuclei of total nuclei (n=3; 1 representative quantified). Statistical analysis was performed using unpaired Student’s *t*-tests. ns, not significant; ****, p<0.0001.

To further delineate the contribution of LGP2 to the sensing of unedited self RNA, we performed a similar experiment in the human cell line HEK293. Knockout of RIG-I, MDA5, or both by CRISPR/Cas9-mediated gene editing was confirmed by immunoblotting and intact type I IFN receptor signaling in the selected clones was verified by monitoring ISG60 expression upon recombinant type I IFN treatment (Fig. 1C). Unexpectedly, siRNA-mediated depletion of ADAR1 did not yield sign of a type I IFN response in parental HEK293 cells or its CRISPR/Cas9-engineered derivatives (Fig. 1D). We noted that these HEK293 cells expressed nearly undetectable levels of LGP2, even after stimulation with recombinant type I IFN (Supplemental Fig. 1A). Importantly, ectopic expression of LGP2 by means of retroviral transduction and stable integration of a FLAG-tagged LGP2-encoding vector (Fig. 1C) enabled type I IFN induction upon siRNA-mediated ADAR1 depletion, as determined by expression of IFN-β and IFIT1 transcripts (Fig. 1D). This was evident in MDA5-sufficient but not in MDA5-deficient cells, confirming that LGP2 and MDA5 were both necessary. As expected, loss of RIG-I did not significantly impact type I IFN induction upon ADAR1 knockdown (Fig. 1D). Of note, increased MDA5 expression, through pre-treatment with recombinant type I IFN, did not bypass the requirement for LGP2 (Supplemental Fig. 2A-B), which suggests that MDA5 levels are not the rate-limiting factor. The ISG signature reached its maximum around 78 hours post-siRNA delivery to LGP2-overexpressing cells (Supplemental Fig. 1B). These observations were confirmed at protein level: ADAR1 depletion upon siRNA treatment led to robust upregulation of the protein ISG60 exclusively in cells that express both MDA5 and LGP2 (Fig. 1E). Moreover, the presence of both LGP2 and MDA5 was required for phosphorylation of IRF3 and STAT1, two key transcription factors that act downstream of MAVS and IFNAR to induce IFN-β and ISG transcription, respectively (Fig. 1E). Finally, nuclear translocation of IRF3, a hallmark of type I IFN induction, only occurred upon expression of LGP2 in ADAR1-depleted cells (Fig. 1F). These observations demonstrate that LGP2 is essential for type I IFN induction and signaling upon ADAR1 depletion.

Previous studies indicated that LGP2 can function as a concentration-dependent biphasic switch that favors MDA5 signaling in response to viral ligands at low concentrations while inhibiting MDA5-dependent responses at high concentrations (Rodriguez *et al*, 2014). However, in our experiments, increasing amounts of an LGP2-encoding plasmid led to a gradual increase in the type I IFN response upon siRNA-mediated ADAR1 depletion without any signs of an inhibitory effect (Supplemental Fig. 1C and 1D) except at very high doses of LGP2, which negatively affect cell viability (not shown). The LGP2-dependent biphasic response previously reported in the context of viral dsRNA sensing is therefore not evident in self RNA sensing.

The absolute requirement for LGP2 in the induction of a type I IFN response following ADAR1 depletion was surprising and distinct from its role in viral dsRNA sensing, where LGP2 potentiates MDA5 signaling but is not strictly required. Indeed, while bona fide LGP2-knockout HEK293 cells failed to induce a type I IFN response upon ADAR1 depletion (Supplemental Fig. 2A-B), they retained the ability to induce a (minimal) type I IFN response upon stimulation with the dsRNA mimic HMW poly(I:C) or RNA isolated from EMCV-infected cells, both of which activate MDA5 (Supplemental Fig. 2E-F). As a control, the siADAR1-induced IFN response was restored in LGP2-knockout cells upon ectopic LGP2 expression (Supplemental Fig. 2C-D). Whether the differential detection of self RNA versus viral RNA by LGP2/MDA5 is caused by a qualitative or quantitative difference, or both, is not clear. Unedited self RNA may either be less abundant in cells or be a less suitable MDA5 ligand (for example because it contains only short stretches of base-paired regions as opposed to long dsRNA found in viral RNA) and therefore it may be more reliant on LGP2 for its detection. Either way, it is evident that the requirement for LGP2 becomes critical in the case of an ‘imperfect’ MDA5 ligand. Altogether, these findings implicate human LGP2 as a key player in the response to unedited self RNA in ADAR1-depleted cells.

### Sensing of unedited self RNA via LGP2 requires RNA binding and ATP hydrolysis

The limited expression of LGP2 and the absence of a type I IFN response upon ADAR1 depletion in wild type HEK293 cells allowed us to create ADAR1 knockout cells through CRISPR/Cas9, without the activation of innate immune pathways that hinder cell proliferation. Two ADAR1-knockout clones were selected that completely lost expression of the ADAR1 p110 and p150 isoform yet remained responsive to type I IFNs, as determined by immunoblotting (Fig. 2A). Genetic loss of ADAR1 did not reveal a type I IFN response until introduction of LGP2 (Fig. 2B-C and Supplemental Fig. 3A-B), in line with our earlier observations using ADAR1 siRNAs. As reported (Pestal *et al*, 2015), the IFN response was largely due to the loss of the p150 isoform, as reconstitution of p150 expression completely blocked the IFN induction in LGP2-expressing ADAR1 knockout cells (Fig. 2B-C and Supplemental Fig. 3A-B). In contrast, overexpression of the p110 isoform reduced, but did not block, this type I IFN response. The reduction can most likely be explained by overexpression of this isoform, which is normally restricted to the nucleus but can “spill” into the cytosol in overexpressing cells. The ADAR1-deficient cells with tunable, LGP2-dependent type I IFN response, provide us therefore with a useful tool to dissect the features of LGP2 and its interaction partners that are required for unedited self RNA sensing.

**Figure 2:**
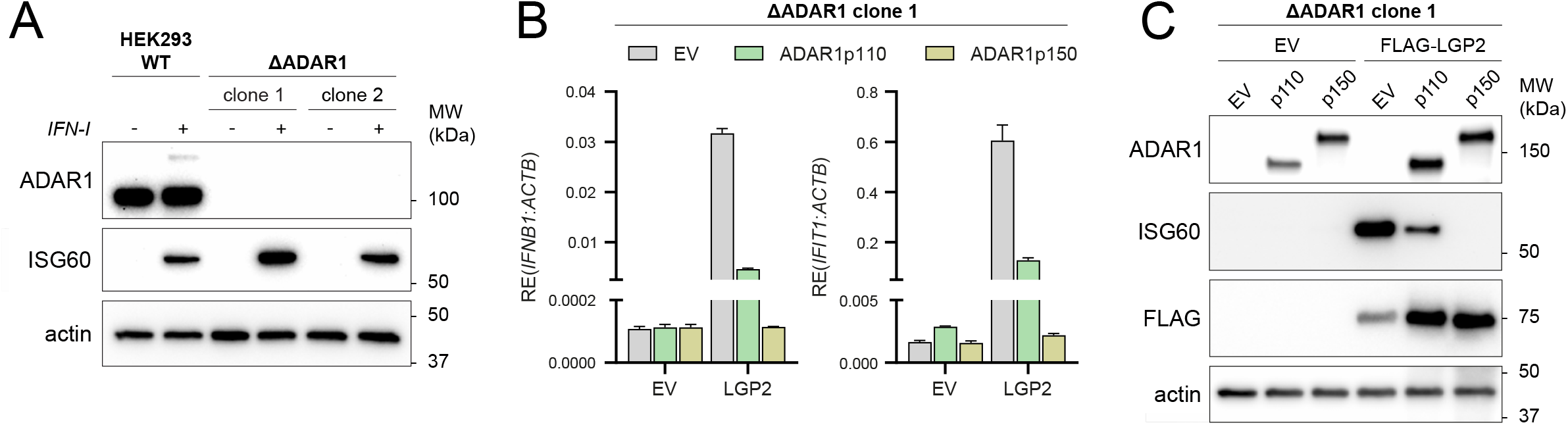
A type I IFN response is unleashed in ADAR1 knockout cells upon expression of LGP2. **A)** HEK293 cells were genetically engineered to knockout ADAR1 using CRISPR/Cas9. Cells were treated for 24h with recombinant type I IFN to upregulate ADAR1 p150 and ISG60 to confirm correct gene editing and type I IFN responsiveness, respectively. Protein lysates were analyzed by SDS-PAGE followed by immunoblotting using the indicated antibodies (n=3). **B)** ADAR1-knockout HEK293 cells (clone 1) were co-transfected with an empty vector (EV) or a FLAG-LGP2-encoding vector (LGP2) combined with a vector encoding GFP-tagged ADAR1 p110 or p150. Cells were harvested 72h post-transfection and the type I IFN response was monitored by RT-qPCR analysis of IFN-β and IFIT1 expression, normalized to ACTB. Data from a representative experiment are shown with mean ± s.d. (n=3). **C)** ADAR1-knockout cells (clone 1) were transfected as in (B). Protein lysates were analyzed by SDS-PAGE followed by immunoblotting using the indicated antibodies (n=3).

The canonical function of LGP2 as an RNA sensor involves RNA binding and ATP hydrolysis while other roles, such as interaction with MAVS and TRAFs, do not (Esser-Nobis *et al*, 2020; Parisien *et al*, 2018). We introduced, by means of lentiviral transduction, a doxycycline-inducible system to stably express FLAG-LGP2 WT or a mutant that completely fails to bind RNA (FLAG-LGP2 K138E/R490E/K634E, denoted as “LGP2 KRK” in figures) in ADAR1 KO cells (Fig. 3A-B). Doxycycline-induced expression of LGP2 WT in ADAR1 KO cells led to robust ISG60 protein (Fig. 3A) and IFN-β and IFIT1 transcript induction (Fig. 3B). In contrast, expression of the LGP2 RNA-binding mutant did not induce a type I IFN response. Consistent with these findings, induction of LGP2 WT, but not the RNA-binding mutant, allowed nuclear translocation of IRF3 (Fig. 3C). These observations indicate that binding to RNA substrates is required for LGP2-dependent type I IFN induction in ADAR1-deficient cells.

**Figure 3:**
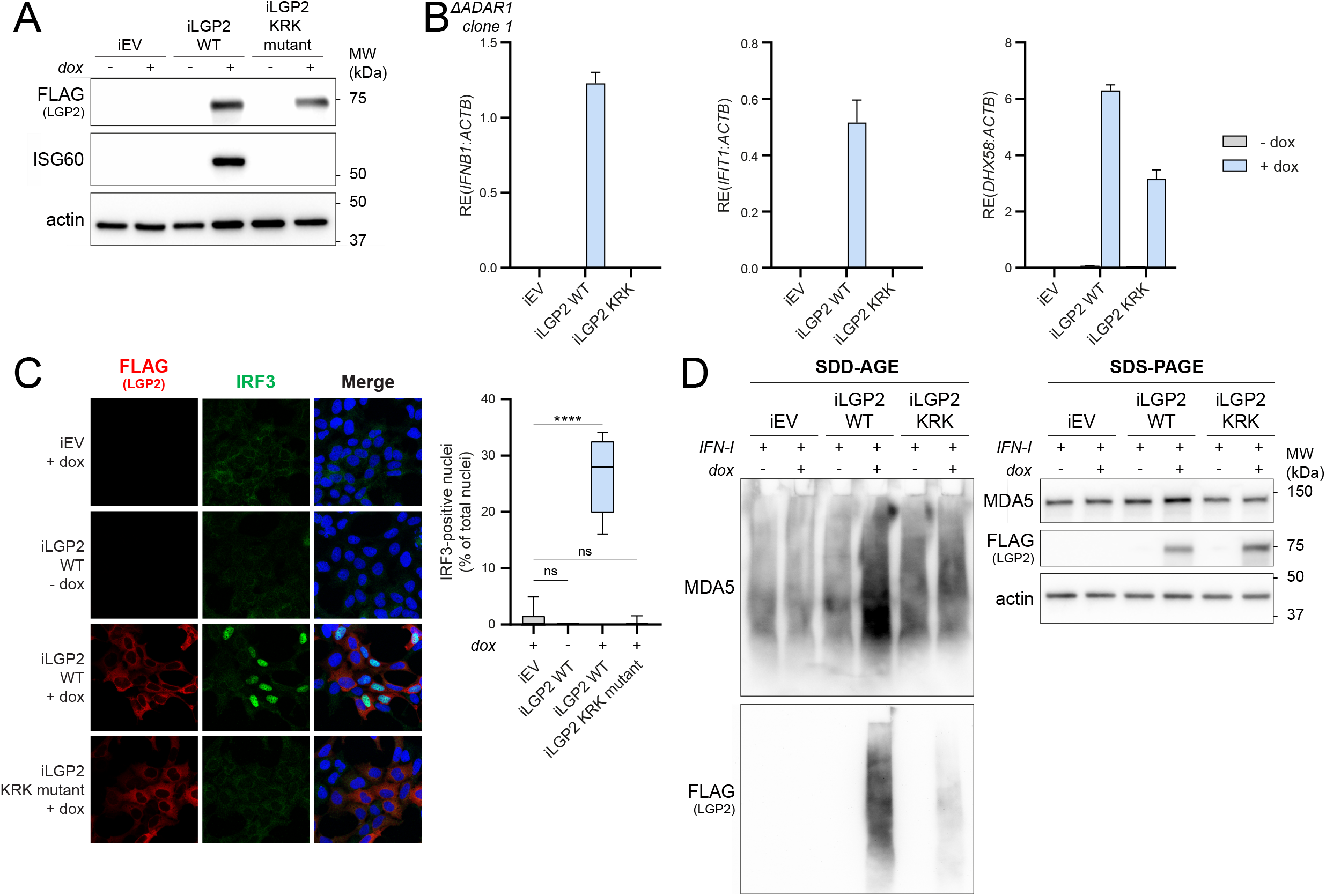
RNA binding by LGP2 is required for receptor oligomerization and type I IFN induction in ADAR1 knockout cells. **A)** ADAR1-knockout HEK293 cells (clone 1) were modified with a lentiviral-based inducible system to express FLAG-LGP2 WT or a FLAG-LGP2 RNA binding mutant (K138E/R490E/K634E, denoted as “KRK mutant”) in a doxycycline-regulated manner. Cells were treated 72h with doxycycline (dox). Protein lysates were analyzed by SDS-PAGE and immunoblotting using the indicated antibodies (n=3). iEV = inducible empty vector; iLGP2 = inducible LGP2. **B)** Cells generated in (A) were treated with doxycycline for 72h to induce LGP2 WT or KRK mutant gene expression. The type I IFN response was monitored by RT-qPCR analysis of IFN-β and IFIT1 transcript expression, normalized to ACTB. Gene induction was monitored by DHX58 (encoding LGP2) transcript expression, normalized to ACTB. Data from a representative experiment are shown with mean ± s.d. (n=3). **C)** Cells generated in (A) were plated on coverslips and treated with or without doxycycline for 72h. Cells were fixed, permeabilized and stained with anti-FLAG and anti-IRF3 antibodies. Total nuclei (>500 nuclei per experimental condition) and IRF3-positive nuclei were counted using semi-automated software analysis and plotted as percentage IRF3-positive nuclei of total nuclei (n=2; 1 representative quantified). Statistical analysis was performed using one-way ANOVA with Sidak’s correction for multiple comparisons. ns, not significant; ****, p<0.0001. **D)** Cells generated in (A) were treated with doxycycline for 72h. During the last 24h, recombinant type I IFN was added to upregulate endogenous MDA5 protein expression. Protein lysates were analyzed by SDD-AGE and SDS-PAGE using the indicated antibodies to determine protein oligomerization and total expression levels, respectively (n=3).

To determine whether LGP2 is required for MDA5 oligomerization in ADAR1-deficient cells, we utilized semi-denaturing detergent agarose gel electrophoresis (SDD-AGE) to monitor MDA5 aggregation. To circumvent discrepancies in MDA5 protein levels across samples (due to its increased expression as an ISG in LGP2-expressing ADAR1 KO cells), we treated cells with recombinant type I IFN to equalize MDA5 expression (Fig. 3D, SDS-PAGE). Doxycycline-inducible expression of LGP2 WT, but not the RNA-binding mutant, revealed MDA5 aggregation in ADAR1 knockout cells (Fig. 3D, SDD-AGE). SDD-AGE further revealed that RNA binding-competent LGP2 oligomerizes in ADAR1 knockout cells (Fig. 3D), consistent with previous studies showing that human and chicken LGP2 itself can form filaments (Bruns *et al*, 2014; Uchikawa *et al*, 2016). These findings place LGP2 at the level of MDA5 oligomerization and activation in the ADAR1-induced type I IFN response.

We transiently expressed various LGP2 truncation mutants and point mutants (Fig. 4A) in ADAR1 knockout cells and observed that, besides RNA binding, full length LGP2 and its ability to hydrolyze ATP are strictly required to sustain a type I IFN response. Expression of the LGP2 N-terminal domain (NTD), C-terminal domain (CTD), or mutation of LGP2 residues that are critical for ATPase activity (K30A) or RNA binding via the LGP2 NTD (K138E/R490E) or CTD (K634E) (Pippig *et al*, 2009; Uchikawa *et al*, 2016; Bruns *et al*, 2013), all abolished IFN induction in ADAR1-deficient cells (Fig. 4B-C).

**Figure 4:**
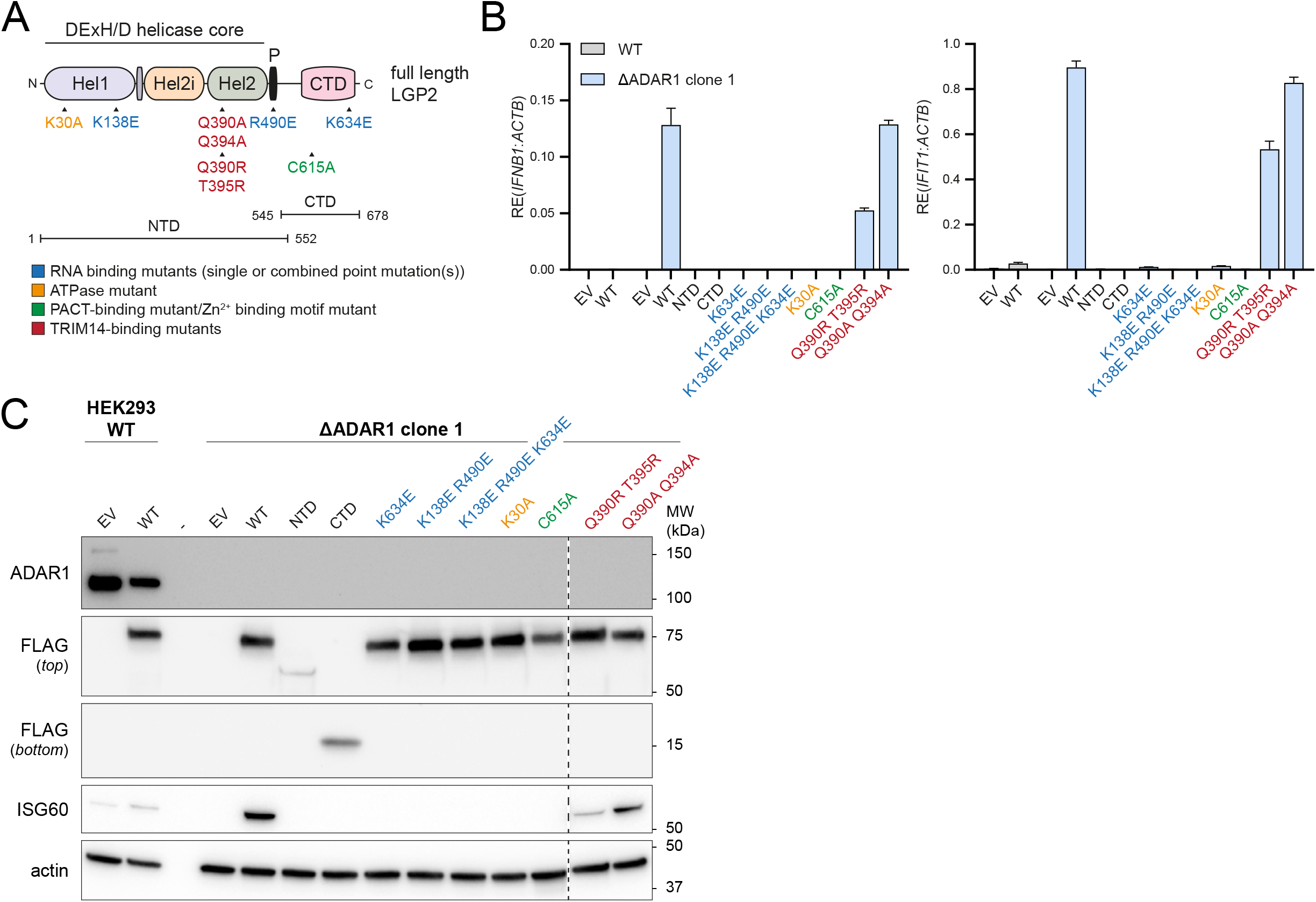
The function of LGP2 in sensing unedited self RNA involves its canonical role as dsRNA sensor that requires RNA binding and ATP hydrolysis. **A)** Schematic illustration of the domain structure of LGP2 and various point mutants and truncation mutants that are used in this study. The N-terminal domain (NTD) of LGP2 is composed of a conserved DExH/D helicase domain, subdivided into the helicase 1 (Hel1), helicase 2 (Hel2) and helicase insertion (Hel2i) domain, and a pincer motif (P). The NTD is followed by a C-terminal domain (CTD), involved in RNA binding. **B)** HEK293 WT or ADAR1-knockout cells (clone 1) were transiently transfected with vectors encoding the indicated truncation or point mutants and harvested 72h post-transfection. The type I IFN response was monitored by RT-qPCR analysis of IFN-β and IFIT1 transcript expression, normalized to ACTB. Data from a representative experiment are shown with mean ± s.d. (n=3). **C)** Cells were treated as in (B). Protein lysates were prepared and analyzed by SDS-PAGE followed by immunoblotting using the indicated antibodies (n=3).

Mutation of a cysteine residue in the C-terminal domain of LGP2 crucial for binding to the dsRNA-binding protein PACT (C615A) (Sanchez David *et al*, 2019) also prevented type I IFN induction, suggesting that PACT, via LGP2, may participate in the IFN response to unedited self RNA (Fig. 4B-C). Of note, C615 is also important for the correct orientation of a Zn^2+^ ion in LGP2 (Pippig *et al*, 2009), hence the role of PACT and/or Zn^2+^ binding will need to be evaluated in further studies.

A recent study identified a biochemical interaction between LGP2 filamentous structures and TRIM14, an unusual member of the TRIM family that lacks a RING domain and does not function as a ubiquitin E3 ligase (Kato *et al*, 2021). Mutations in the α3 helix of the Hel2 domain of LGP2 (Q390R/T395R or Q390A/Q394A) strongly decreased the interaction between the Hel2i-Hel2 domain of LGP2 and TRIM14 (Kato *et al*, 2021). We found that the Q390A/Q394A mutation did not impact on the ability of LGP2 to induce a type I IFN response in ADAR1 knockout cells, while the Q390R/T395R mutation reduced, but not abolished, type I IFN induction (Fig. 4B-C). Whether TRIM14 plays a more pronounced role in type I IFN induction upon picornavirus infection or perhaps regulates alternative, non-canonical functions of LGP2, needs to be further explored in a TRIM14-deficient setting.

We conclude that, besides RNA binding, ATP hydrolysis is strictly required for LGP2 to mediate type I IFN induction in ADAR1-deficient cells. This suggests that the function of LGP2 in inflammation in ADAR1 deficiency involves its canonical role as RNA sensor rather than an ‘indirect’ role, e.g., via its interaction with MAVS or TRAFs. In addition, the binding of LGP2 to PACT or a Zn^2+^ ion is important for type I IFN induction in ADAR1-deficient cells.

### LGP2 is required for growth retardation of tumor cells and cell-intrinsic inflammation upon loss of ADAR1, which is potentiated by epigenetic therapy

Recent studies have placed ADAR1 in the spotlight as an attractive novel drug target to enhance anti-tumor immunity (Bhate *et al*, 2019). To explore the relationship between LGP2 and ADAR1 and its prognostic value for overall patient survival, we performed *in silico* analysis of ADAR1 and LGP2 (encoded by *ADAR* and *DHX58*, respectively) mRNA expression in multiple cancer types using data from The Cancer Genome Atlas (TCGA). We hypothesized that patients with high *DHX58* expression (*DHX58^high^*) would have improved survival compared to patients with low *DHX58* expression (*DHX58^low^*) in patients with low *ADAR* expression (*ADAR^low^*). Patients were stratified into four groups according to *ADAR* and *DHX58* transcript levels using median cut-offs. Consistent with the hypothesis, *ADAR^low^DHX58^high^* patients had improved overall survival compared to *ADAR^low^DHX58^low^* patients (Fig. 5A and Supplemental Fig. 4). In contrast, no difference in survival was found in *ADAR^high^* cancer patients stratified based on *DHX58* levels. This observation was most striking in bladder cancer (BLCA), sarcoma (SARC) and breast cancer (BRCA) (Fig. 5A). Thus, LGP2 mRNA abundance is linked to improved outcomes specifically in patients with low levels of ADAR1 across multiple malignancies.

**Figure 5:**
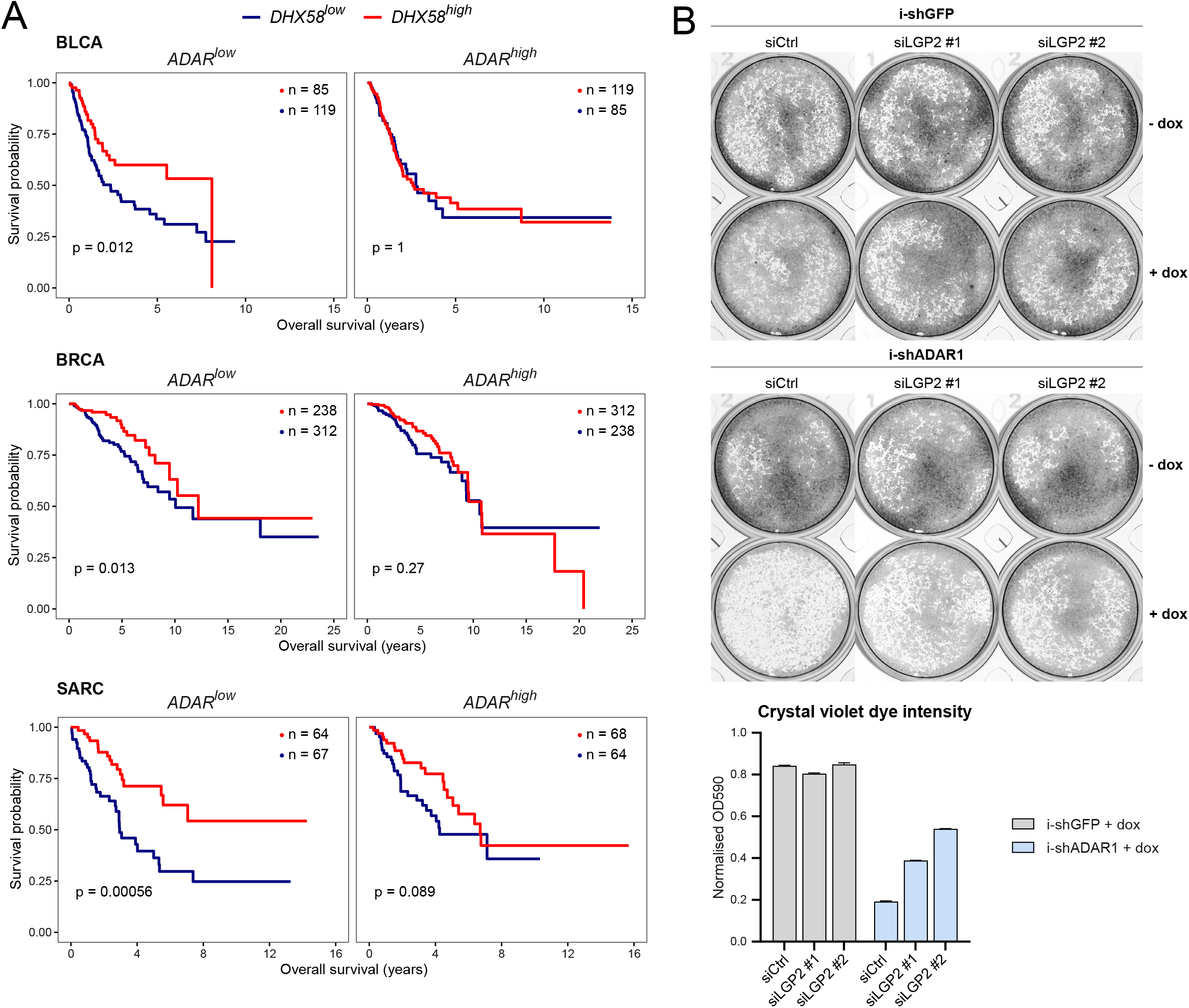
Reduced tumor cell growth upon loss of ADAR1 is dependent on LGP2. **A)** Kaplan-Meier plots showing overall survival of *ADAR*^low^ and *ADAR*^high^ patients stratified by *DHX58* levels in bladder cancer (BLCA; n=407), breast cancer (BRCA; n=1099) and sarcoma (SARC; n=263) TCGA datasets. Median cut-offs were used for patient stratification and logrank test P values are shown. **B)** CAL27 cells transduced with doxycycline-inducible shRNAs targeting ADAR1 or GFP (negative control) were treated with doxycycline and transfected with two independent siRNAs targeting LGP2 (siLGP2 #1 or #2) or a control siRNA (siCtrl). 120h post-transfection, cells were fixed, stained with crystal violet and imaged. Crystal violet was extracted from stained cells and the dye intensity was quantified using a colorimetric assay (OD_590_). OD_590_ values of doxycycline-treated cells were normalized to the OD_590_ values of untreated cells. Quantification data of a representative experiment are shown (n=2).

Whether the above observations indicate a functional relationship between ADAR1 and LGP2 in tumors, or are merely a consequence of increased ISG expression in tumors with low ADAR1 expression, cannot be determined through bioinformatic analysis. To explore this experimentally, we used doxycycline-inducible shRNA-mediated knockdown of ADAR1 in a human oral squamous cell carcinoma cell line (CAL27), which was previously shown to be sensitive to ADAR1 loss (Liu *et al*, 2019) and that upregulates LGP2 upon type I IFN treatment (Supplemental Fig. 5A). As predicted, knockdown of ADAR1 inhibited cell proliferation of CAL27 cells, whereas this was not the case upon expression of a control shRNA targeting GFP (Fig. 5B). Importantly, simultaneous knockdown of LGP2 using two independent siRNAs resulted in a partial rescue of cell growth after ADAR1 knockdown (Fig. 5B). ADAR1 and LGP2 knockdown efficiencies were validated by RT-qPCR (Supplemental Fig. 5B). The partial rescue of ADAR1-mediated growth retardation is likely attributable to the incomplete knockdown of LGP2 as well as the activation of PKR upon ADAR1 loss, as reported (Chung *et al*, 2018; Liu *et al*, 2019). We conclude that loss of LGP2 in part suppresses ADAR1 shRNA-induced growth retardation.

Treatment of cancer cells with DNA methyltransferase inhibitors (DNMTis) activates RNA sensors, including MDA5 and the endosomal dsRNA sensor Toll-like receptor 3 (TLR3) (Roulois *et al*, 2015; Chiappinelli *et al*, 2015). Moreover, the combined treatment of patient-derived colorectal cancer cell lines with the DNMTi 5-aza-2’-deoxycytidine (5-AZA-CdR) together with shRNA-mediated depletion of ADAR1 induces an MDA5-dependent type I IFN response through the increased expression of IR-Alu elements that are no longer edited (Mehdipour *et al*, 2020). To test whether this involves LGP2, we used siRNAs to deplete ADAR1, either alone or in combination with an siRNA targeting LGP2, in human colorectal adenocarcinoma cells (HT29). As expected, loss of ADAR1 triggered a strong type I IFN response in these cells, as determined by expression of IFN-β and two ISGs (IFIT1 and ISG15), but this was completely blocked upon simultaneous depletion of LGP2 (Supplemental Fig. 5C). The siADAR1-mediated IFN response was further enhanced upon 5-AZA-CdR treatment of HT29 cells (Fig. 6A), as previously reported (Mehdipour *et al*, 2020). Notably, the synergistic effect between ADAR1 depletion and 5-AZA-CdR treatment was also strictly dependent on LGP2 (Fig. 6A). In contrast, the elevated type I IFN response induced by treatment with 5-AZA-CdR alone was less dependent on LGP2, which indicates that some stimulatory RNAs that are demethylated and expressed upon 5-AZA-CdR treatment *per se* partially escape recognition by LGP2, likely through ADAR1-mediated RNA editing. Knockdown efficiencies of ADAR1 and LGP2 were determined by RT-qPCR analysis (Supplemental Fig. 5C). The above observations were confirmed in another human colorectal carcinoma cell line (LIM1215) (Fig. 6B and Supplemental Fig. 5D). To further explore the LGP2-dependent synergy between ADAR1 depletion and anti-cancer therapies, we treated cells with ADAR1-targeting siRNAs in the presence or absence of the CDK4/6 inhibitor palbociclib, which also upregulates levels of endogenous stimulatory RNAs (Goel *et al*, 2017). Combined treatment with ADAR1 siRNAs and palbociclib caused a synergistic upregulation of IFN-β and ISGs, as described (Mehdipour *et al*, 2020), which was also strictly dependent on LGP2 (Fig. 6C). We conclude that LGP2 is essential for the enhanced inflammatory response upon combined epigenetic therapy and ADAR1 depletion in tumor cells.

**Figure 6:**
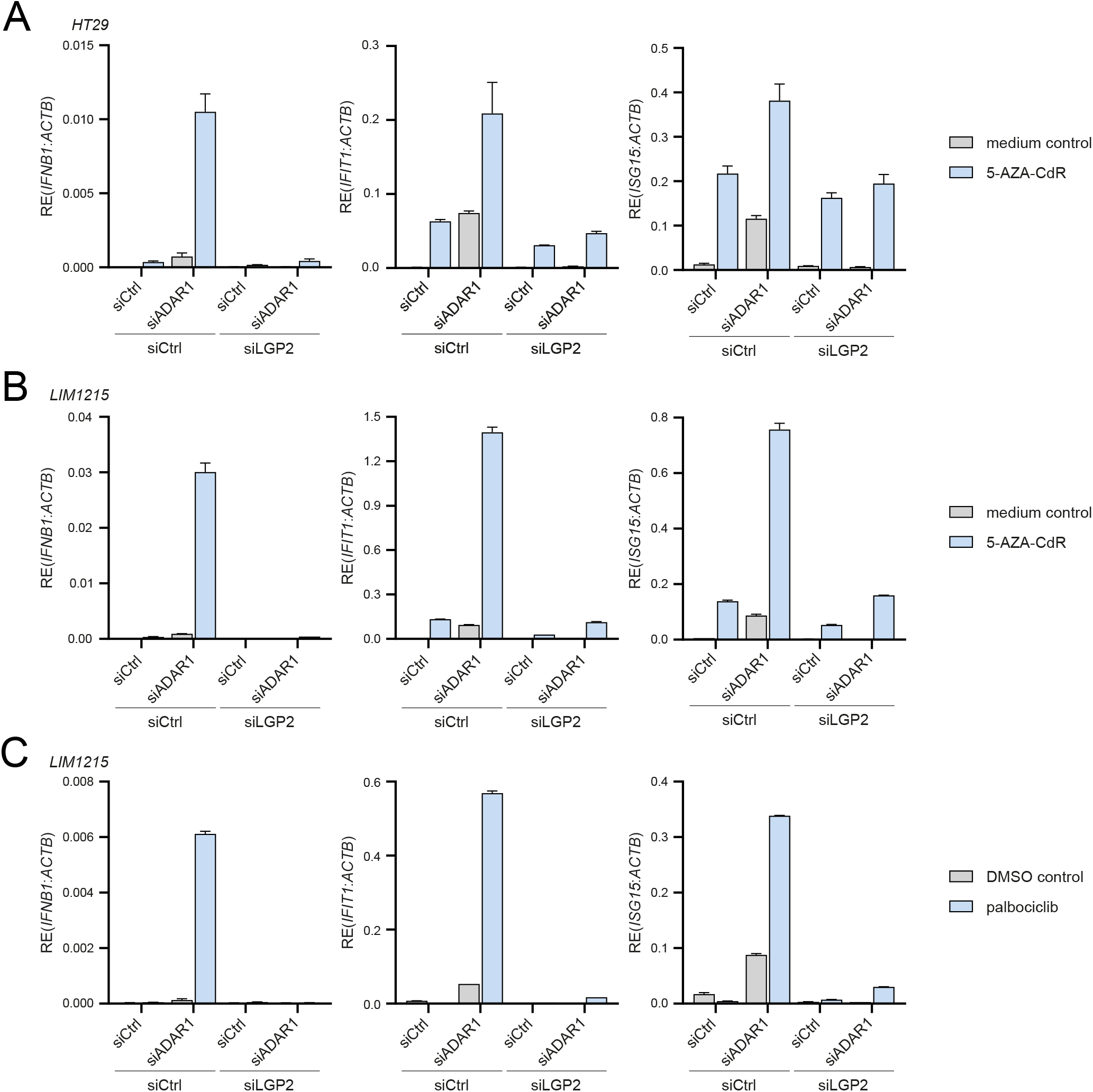
Depletion of ADAR1 in combination with epigenetic repressors trigger type I IFN induction in an LGP2-dependent manner. **A)** HT29 cells were treated with or without 300 nM 5-AZA-CdR for 2 days and subsequently washed and transfected with the indicated siRNAs. Cells were harvested 72h post-transfection and the type I IFN response was analyzed by RT-qPCR analysis of IFN-β, IFIT1, and ISG15 transcript expression, normalized to ACTB. Data from a representative experiment are shown with mean ± s.d. (n=2). **B)** LIM1215 cells were treated with or without 300 nM 5-AZA-CdR for 2 days and subsequently washed and transfected with the indicated siRNAs. Cells were harvested 72h post-transfection and the type I IFN response was analyzed as in (C) (n=2). **C)** LIM1215 cells were treated with 250 nM palbociclib or a DMSO control for 7 days. Three days after treatment initiation, cells were transfected with the indicated siRNAs and cultured for an additional 72h. The type I IFN response was monitored as in (C) (n=2).

## Discussion

Nucleic acid sensors continuously survey their environment for the presence of nucleic acid structures that are commonly found on viral RNA. Various cellular mechanisms allow the discrimination between viral (non-self) nucleic acids and cellular (self) DNA or RNA. These mechanisms, however, are not foolproof. Loss of ADAR1-dependent RNA editing causes unwanted recognition of self RNA and consequently inadvertent innate immune activation and severe pathology (Rodero & Crow, 2016; Rice *et al*, 2012, 2017; Livingston & Crow, 2016). Various nucleic acid sensors, including PKR, OAS/RNase L and MDA5, sense unedited self RNA and cause translational shutdown, cell death and autoinflammation, marked by the production of type I IFNs (Quin *et al*, 2021). Here we demonstrate that the RNA helicase LGP2 is indispensable for type I IFN induction in ADAR1-deficient human cells. We further demonstrate that this involves the canonical role of LGP2 as a sensor of base-paired RNA, which requires ATP hydrolysis and intact RNA binding sites and enables MDA5 oligomerization. We extend our findings to multiple cancer cell lines (THP-1, CAL27, HT29 and LIM1215) and demonstrate that the sensitivity of tumor cells to ADAR1 loss requires the presence of LGP2. Moreover, the previously reported synergistic effects of ADAR1 depletion and epigenetic therapy on the intrinsic type I IFN response in cancer cells are also strictly dependent on LGP2. These findings have several important implications, which are discussed below.

The key role of LGP2 in the sensing of unedited self RNA sheds new light on the search for the, largely elusive, stimulatory RNAs in ADAR1 deficiency. Multiple studies have defined the ADAR1 ‘editome’ (Ramaswami & Li, 2016). The vast majority of A-to-I editing sites is found in or near Alu elements, and a small proportion is found in other mobile repeat elements, such as long interspersed nuclear elements (LINEs) and endogenous retroviruses (ERVs) (Ramaswami *et al*, 2012). Alu elements are mostly embedded within 3’UTRs or introns of Pol II transcripts, while few are transcribed as individual units by Pol III (Chung *et al*, 2018; Deininger, 2011). The repetitive nature of mobile elements, especially when found in close proximity to each other and in inverted orientation, increases the risk of forming endogenous base-paired structures that may activate innate immune pathways (Eisenberg & Levanon, 2018). Editing of repeat elements by ADAR1 may decrease the high degree of complementarity in such RNAs and minimize accidental innate immune activation. An elegant study indeed demonstrated that, *in vitro,* recombinant MDA5 preferentially binds unedited IR-Alu elements amongst total cytosolic RNA extracted from ADAR1-deficient cells (Ahmad *et al*, 2018). However, editing levels are often low within base-paired stem regions of Alu and IR-Alu elements while being more frequent within predicted single stranded regions (Chung *et al*, 2018). In addition, endogenous base-paired RNAs with near-perfect complementarity are scarce amongst mRNAs (but not pre-mRNAs) (Barak *et al*, 2020). In mice, transcriptomic analysis of ADAR p110/ADAR2-deficient brain tissue revealed as few as 36 ADAR1 p150-specific editing sites (Kim *et al*, 2021). Thus, potentially, only few RNAs become truly stimulatory when no longer edited in intact cells or tissues. The stimulatory potential of endogenous RNA also depends on its conformation. RNA that adopts a non-canonical Z-conformation has increased immunostimulatory potential. The latter is reduced upon binding and editing by ADAR1 p150, which contains a Z-nucleic acid (Zα) binding domain (de Reuver *et al*, 2021; Tang *et al*, 2021; Nakahama *et al*, 2021; Maurano *et al*, 2021; Zillinger & Bartok, 2021). In mice, various mutations in the Zα domain of ADAR1 result in postnatal growth retardation and mortality as well as an increased type I IFN signature (de Reuver *et al*, 2021; Maurano *et al*, 2021; Nakahama *et al*, 2021; Tang *et al*, 2021). Transcripts that are prone to adopt a Z-conformation, such as those with purine-pyrimidine repeats (Koeris *et al*, 2005), can therefore be immunostimulatory and this can be reduced by A-to-I editing. The precise identity and features of immunogenic RNAs remains unresolved. Protein-RNA interaction studies, such as individual-nucleotide resolution UV-crosslinking and immunoprecipitation (iCLIP), may help to identify RNAs that directly engage dsRNA sensors. The role of LGP2 in unedited self RNA sensing opens up the possibility to retrieve immunogenic RNAs via their association with LGP2 in ADAR1-deficient cells, as has been done in the context of viral infection (Deddouche *et al*, 2014). Our observation that type I IFN induction upon loss of ADAR1 is strictly dependent on LGP2, which favors the formation of short RNA filaments (Bruns *et al*, 2014), hints at the possibility that the stimulatory RNA in ADAR1 deficiency is shorter than ‘classical’ MDA5 substrates, which tend to be long complex dsRNAs of > 1000/2000 nt in length (Pichlmair *et al*, 2009; Kato *et al*, 2021).

Besides ADAR1, eight disease-causative mutations have been identified in AGS, including gain-of-function mutations in *IFIH1* (encoding MDA5), all of which cause an elevated type I IFN signature (Livingston & Crow, 2016; Rodero & Crow, 2016; Uggenti *et al*, 2019, 2020). Currently there is no licensed therapy for treatment of AGS or related interferonopathies (Crow *et al*, 2020). A few individual reports describe encouraging clinical improvements upon treatment of a small number of patients with the JAK1/2 inhibitor ruxolitinib, which blocks signaling downstream of the IFNAR (Crow *et al*, 2020). However, JAK1/2 inhibition is rather non-specific and will block pathways beyond type I IFN signaling. In addition, transcriptional activity of IRF3 and NF-*κ*B leads to upregulation of other genes and cytokines, besides type I IFNs, which perhaps contribute to pathology as well (Andersen *et al*, 2008; Rehwinkel & Gack, 2020). Compounds that target the upstream nucleic acid sensing machinery, such as LGP2, may therefore have therapeutic value. In contrast to ruxolitinib, inhibition of LGP2 will prevent IRF3 and NF-*κ*B activation while minimally impacting on viral nucleic acid sensing via RIG-I and DNA sensors, leaving patients less prone to a wide spectrum of viral infections.

Our findings on the essential role of LGP2 in type I IFN induction caused by ADAR1 dysfunction are strengthened by a recent report in which *Adar^P195A/p150-^* mice, which bear a mutation in the Zα domain of ADAR1 p150 (P195A) paired with a null allele of *Adar* (mimicking the most common *ADAR* mutation in AGS at P193), were intercrossed with various knockout models, including *Dhx58^-/-^* mice (Maurano *et al*, 2021). LGP2 deficiency rescued the postnatal mortality of *Adar^P195A/p150-^* mice and abolished the type I IFN signature. Thus, LGP2 is an essential effector molecule of ADAR1-driven disease in both mice and humans. Loss of PKR also rescued the mortality of *ADAR^P195A/p150-^* mice, yet the type I IFN signature remained elevated (Maurano *et al*, 2021), consistent with previous literature showing that activation of PKR is largely responsible for ADAR1-associated translational shutdown, cell death and pathology but not IFN-driven inflammation (Chung *et al*, 2018).

In contrast to its disease-causing role in AGS, ADAR1 is an exciting new immuno-oncology target and several *in vitro* and *in vivo* studies have highlighted that its deletion increases tumor cell lethality and renders tumors more vulnerable to immunotherapy (Gannon *et al*, 2018; Liu *et al*, 2019; Ishizuka *et al*, 2019). We observed that depletion of ADAR1 in multiple tumor cell lines triggered a type I IFN response in an LGP2-dependent manner. Moreover, the reported synergy between ADAR1 deletion and epigenetic therapy was completely dependent on the expression of LGP2. Finally, LGP2 was required for delayed cell growth upon ADAR1 knockdown. Altogether this demonstrates that LGP2 is a hitherto overlooked, yet essential, player when targeting ADAR1. It also predicts that LGP2-sufficient tumors are more likely to respond to ADAR1-directed therapies than LGP2-deficient tumors. Indeed, across multiple human tumor types, patient stratification based on ADAR1 and LGP2 transcript levels revealed that patients with high LGP2 and concomitant low ADAR1 levels had improved survival. The relationship between ADAR1 and LGP2 and its impact on tumor growth, the intra-tumoral inflammatory response, and anti-tumor immunity will need to be further evaluated in *in vivo* models.

Collectively, our data identify LGP2 as an important sensor of endogenous stimulatory RNA and as an essential player in autoinflammation driven by ADAR1 dysfunction with important implications for treatment of type I interferonopathies as well as for potential ADAR1-directed cancer therapy.

## Acknowledgements

We are grateful to Dr. Maaike Ressing and Dr. Jan Rehwinkel for critical reading of this manuscript. We thank Dr. Jannie Borst and all members of our laboratory for helpful discussions and suggestions. This work was supported by a fellowship from the Leiden University Medical Centre and a research grant from the Institute for Chemical Immunology (ICI-00203), which is funded by a Gravitation project from the Netherlands Organization for Scientific Research (NWO). CPB and SZ are supported by the NIHR Manchester Biomedical Research Centre and by a Cancer Research UK Manchester Institute Award (A19258). CRS is supported by The Francis Crick Institute, which receives core funding from Cancer Research UK (FC001136), the UK Medical Research Council (FC001136), and the Wellcome Trust (FC001136), by ERC Advanced Investigator Grants (AdG 268670, 786674) by a Wellcome Investigator Award (WT106973MA), and by a prize from the Louis-Jeantet Foundation. This research was funded in whole, or in part, by the Wellcome Trust (grants FC001136, WT106973MA). For the purpose of Open Access, the author has applied a CC BY public copyright license to any Author Accepted Manuscript version arising from this submission.

## Author Contributions

JES, TO, and AGV designed experiments and analyzed data. JES, TO, and AGV conducted experiments with assistance from LTH and DG. CB and SZ performed the TCGA bioinformatic analysis. CRS provided advice and sponsored the initial stages of the project. AGV supervised the project. AGV wrote the manuscript with assistance from JES and TO. All authors reviewed the manuscript.

---

**Supplemental Figure 1, related to Figure 1:**
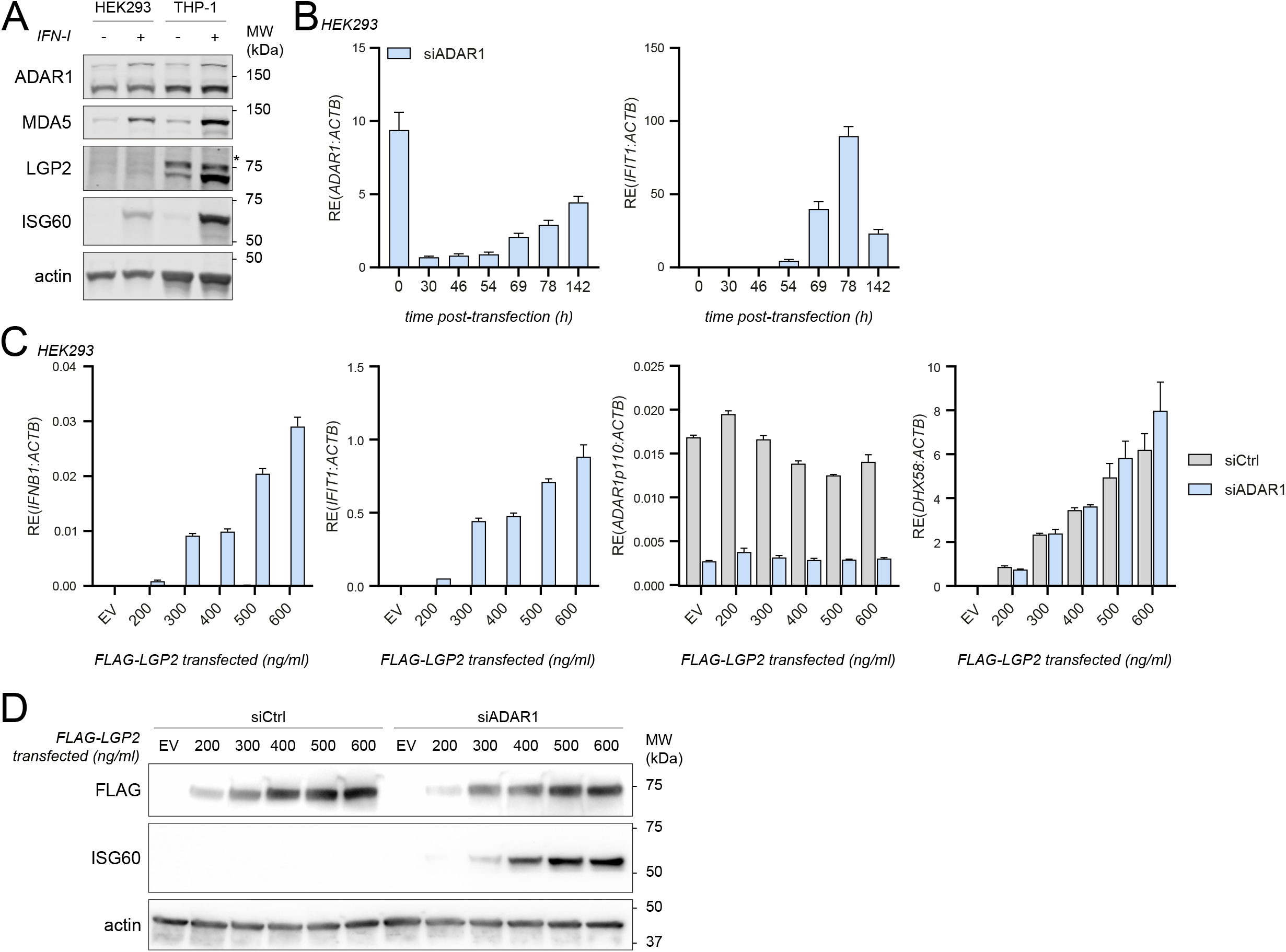
LGP2 is required for the induction of a type I IFN response upon siRNA-mediated depletion of ADAR1 in HEK293. **A)** Expression level and type I IFN-inducibility of relevant proteins in HEK293 and THP-1. HEK293 and PMA-differentiated THP-1 cells were treated with or without recombinant type I IFN. Protein lysates were analyzed by SDS-PAGE followed by immunoblotting with the indicated antibodies (n=3). *, non-specific band. **B)** Kinetics of siRNA-mediated depletion of ADAR1 and induction of the type I IFN response in HEK293. HEK293 cells stably expressing FLAG-LGP2 were transfected with an siRNA targeting ADAR1 (siADAR1) and harvested at the indicated time points post-transfection. ADAR1 knockdown and IFIT1 upregulation were monitored by RT-qPCR analysis (using Taqman probes) and normalized to ACTB. Data from one experiment is shown with mean ± s.d.. **C)** HEK293 WT cells were transfected with siADAR1 or a control siRNA (siCtrl) and 8h later with increasing amounts of a vector encoding FLAG-LGP2. As a control, cells were transfected with 250 ng of an empty vector (EV). Cells were harvested 80h post siRNA transfection. RT-qPCR analysis was used to monitor ADAR1 knockdown, LGP2 (*DHX58*) expression, and IFN-β and IFIT1 expression. All transcripts were normalized to ACTB. Data from a representative experiment is shown with mean ± s.d. (n=2). **D)** Cells were treated as in (C). Protein lysates were prepared and analyzed by SDS-PAGE followed by immunoblotting using the indicated antibodies.

**Supplemental Figure 2, related to Figure 1:**
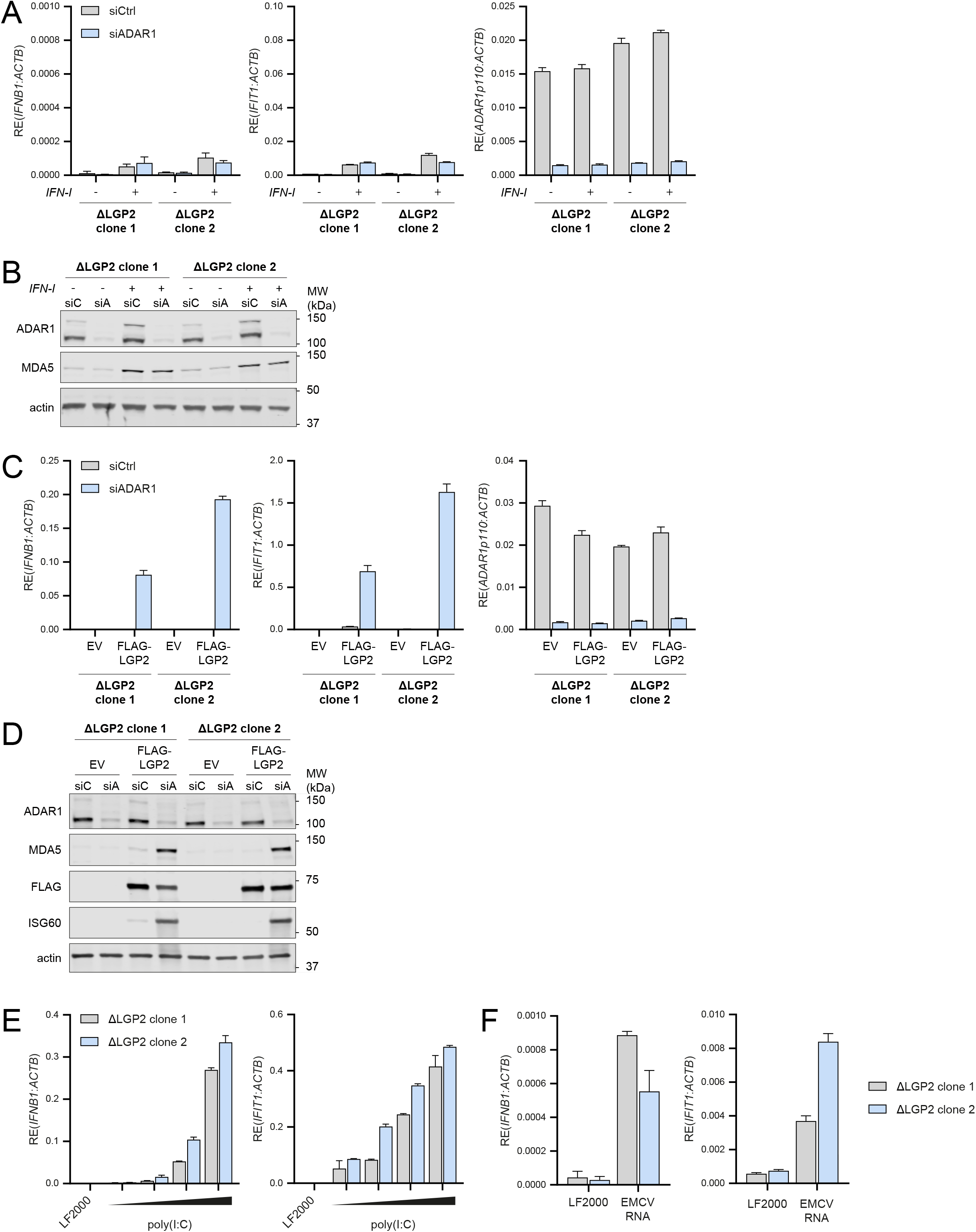
LGP2-deficient cells fail to sense unedited self RNAs, yet maintain the ability to detect viral dsRNAs. **A)** LGP2-knockout HEK293 cells (clones 1 and 2) were transfected with an siRNA targeting ADAR1 (siADAR1) or a control siRNA (siCtrl) and were treated 8h later with recombinant type I IFN to upregulate RLR expression. Cells were harvested 80h post siRNA transfection and RT-qPCR analysis was used to monitor IFN-β and IFIT1 expression and ADAR1 knockdown. All transcripts were normalized to ACTB. Data from a representative experiment is shown with mean ± s.d. (n=3). **B)** Cells were treated as in (A). Protein lysates were prepared 80h post siRNA transfection, followed by SDS-PAGE and immunoblotting using the indicated antibodies (n=3). siC = siCtrl; siA = siADAR1. **C)** LGP2-knockout HEK293 cells (clones 1 and 2) were transfected with siADAR1 or siCtrl and 8h later with a vector encoding FLAG-LGP2 or an empty vector (EV). Cells were harvested 80h post siRNA transfection and RT-qPCR analysis was used to monitor IFN-β and IFIT1 expression and ADAR1 knockdown. All transcripts were normalized to ACTB. Data from a representative experiment is shown with mean ± s.d. (n=2). **D)** Cells were treated as in (C). Protein lysates were prepared 80h post siRNA transfection, followed by SDS-PAGE and immunoblotting using the indicated antibodies (n=2). **E and F)** LGP2-knockout HEK293 cells (clones 1 and 2) were transfected with transfection reagent only (LF2000), increasing amounts (56, 112, 225, 450 ng) of poly(I:C) (E) or 450 ng of RNA isolated from HEK293 cells infected with EMCV in the presence of ribavirin (F). Cells were harvested 16h post-transfection and RT-qPCR analysis was used to monitor IFN-β and IFIT1 expression. All transcripts were normalized to ACTB. Data from a representative experiment is shown with mean ± s.d. (n=2).

**Supplemental Figure 3, related to Figure 2:**
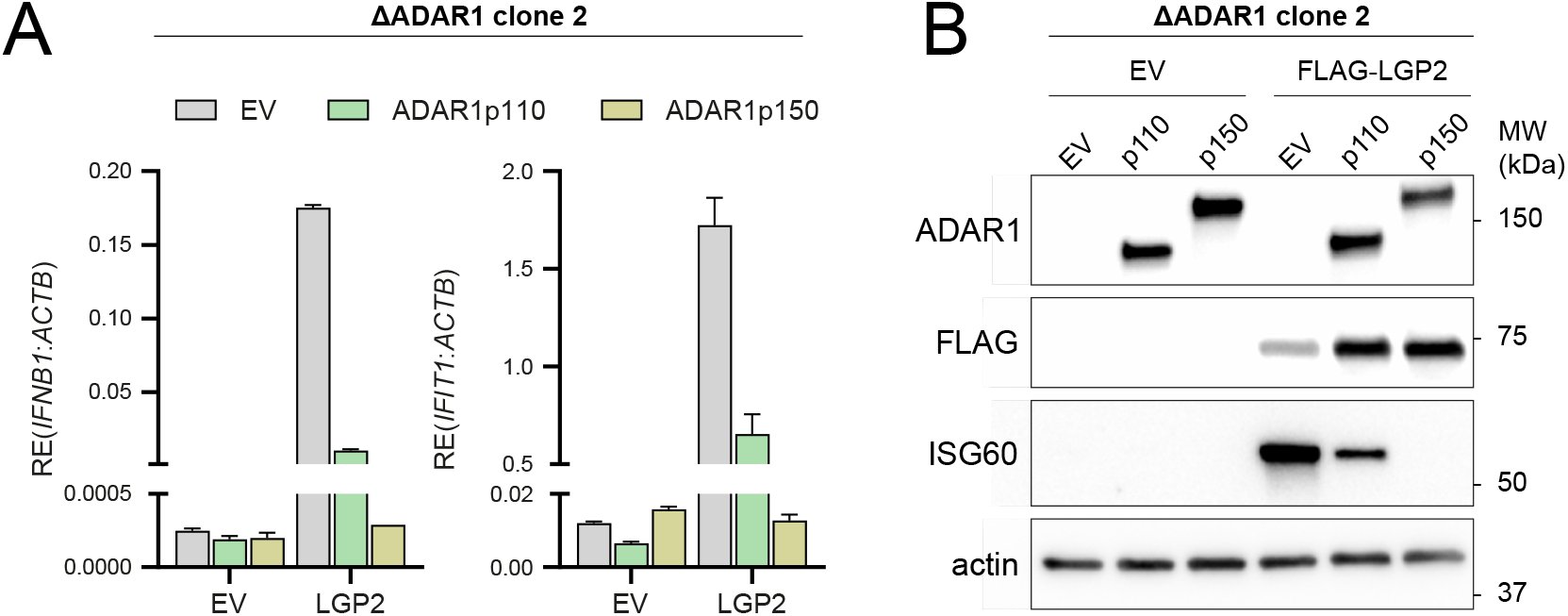
A type I IFN response is unleashed in ADAR1 knockout cells upon expression of LGP2. **A)** ADAR1-knockout HEK293 cells (clone 2) were co-transfected with an empty vector (EV) or a FLAG-LGP2-encoding vector (LGP2) combined with a vector encoding GFP-tagged ADAR1 p110 or p150. Cells were harvested 48h post-transfection and the type I IFN response was monitored by measuring IFN-β and IFIT1 transcript expression, relative to ACTB expression, by RT-qPCR. Data from a representative experiment are shown with mean ± s.d. (n=3). **B)** ADAR1-knockout HEK293 cells (clone 2) were transfected as in (A). Protein lysates were analyzed by SDS-PAGE followed by immunoblotting using the indicated antibodies (n=3).

**Supplemental Figure 4, related to Figure 5:**
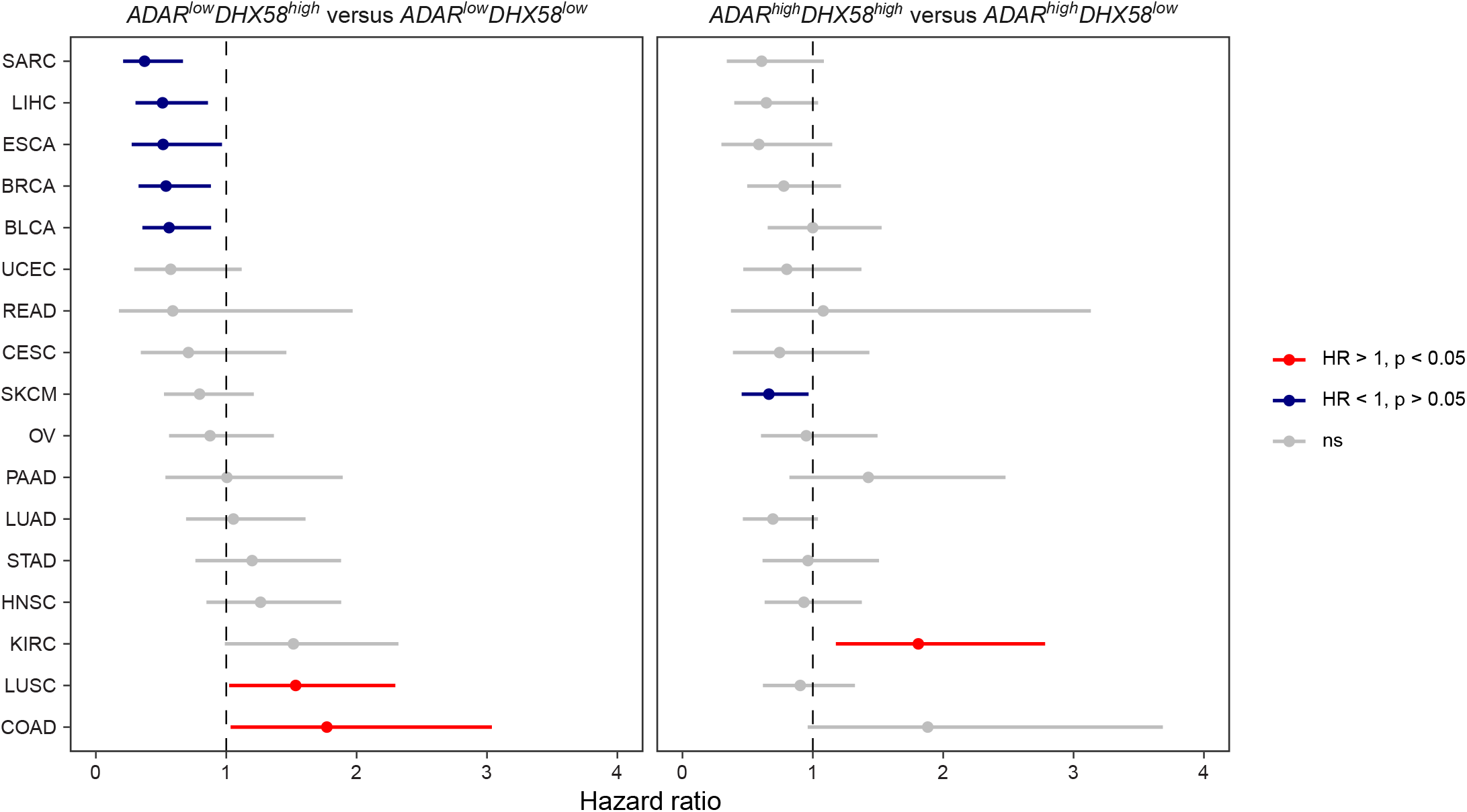
*ADAR*^low^ patients with concomitant *DHX58*^high^ expression have prolonged survival across multiple cancer types. Hazard ratios and 95% confidence intervals from univariate Cox regression models for *DHX58* stratification in *ADAR*^low^ (left panel) and *ADAR*^high^ (right panel) patients from 17 TCGA datasets (sarcoma = SARC, liver = LIHC, esophageal = ESCA, breast = BRCA, bladder = BLCA, endometrial = UCEC, rectal = READ, cervical = CESC, melanoma = SKCM, ovarian = OV, pancreas = PAAD, lung adenocarcinoma = LUAD, stomach = STAD, head and neck = HNSC, clear cell renal cell carcinoma = KIRC, lung squamous = LUSC, colon = COAD). Median cut-off values for both *ADAR* and *DHX58* were used for patient stratification.

**Supplemental Figure 5, related to Figure 5 and 6:**
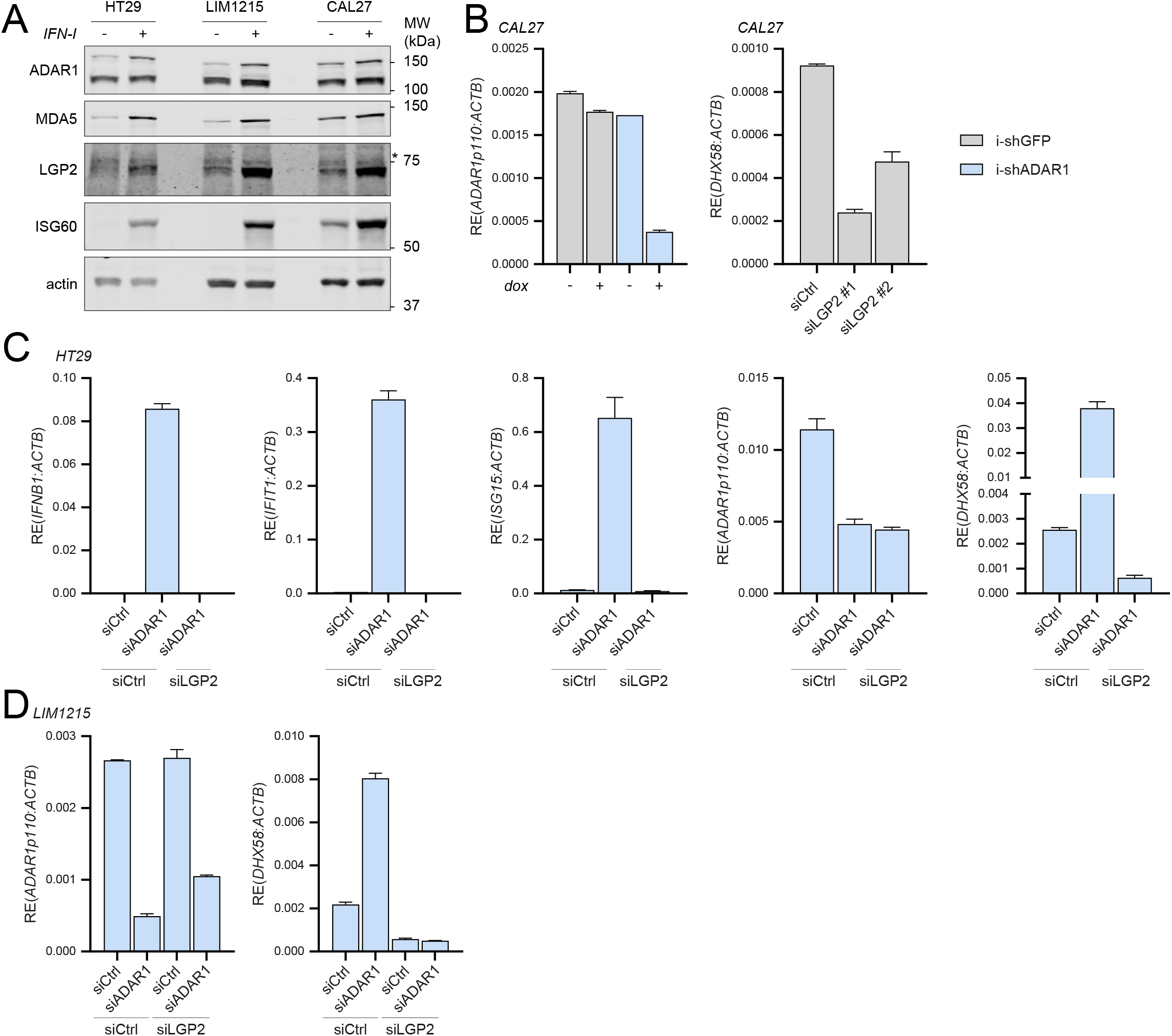
Type I IFN responsiveness and siADAR1-dependent type I IFN induction in various cancer cell lines. **A)** Endogenous expression level and type I IFN inducibility of relevant proteins in HT29, LIM1215, and CAL27 cells. Cells were treated with or without recombinant type I IFN. Protein lysates were analyzed by SDS-PAGE followed by immunoblotting with the indicated antibodies. **B)** Knockdown efficiency of ADAR1 and LGP2 upon doxycycline treatment or siRNA transfection, respectively. ADAR1 and LGP2 (*DHX58*) transcript levels were measured by RT-qPCR and plotted as mean ± s.d. **C)** HT29 cells were transfected with the indicated siRNAs. Cells were harvested 72h post-transfection and RT-qPCR analysis was used to monitor the type I IFN response (IFN-β, IFIT1, and ISG15 transcripts) and knockdown efficiency of ADAR1 and LGP2 (*DHX58*). All transcripts were normalized to ACTB. Data from a representative experiment are shown with mean ± s.d. (n=2). **D)** LIM1215 cells were transfected with the indicated siRNAs. Cells were harvested 72h post-transfection and knockdown efficiency of ADAR1 and LGP2 (*DHX58*) was analyzed as in (C) (n=2).

